# The human complement factor B-C3b complex: Investigation of the interaction using C3b bound to thiol-Sepharose

**DOI:** 10.1101/2024.01.16.575844

**Authors:** Samantha Williams, Kirsten Pondman, Praveen M. Varghese, Ahmad Al Aiyan, Uday Kishore, Robert B. Sim

## Abstract

Factor B, a serine protease proenzyme and part of the complement system, binds to other complement proteins such as C3b and properdin. While it is known to be activated by factor D, other interactions that interfere with the binding between Factor B and C3b are less well-understood. We attached C3b to a thiol Sepharose via its free SH group and conducted a competition assay employing ^125^I-labelled factor B to study competitive binding. Two anti-C3d monoclonal antibodies (F49b and 4C2) were found to partially inhibit the binding of Factor B to C3b. The inhibitory effects of these monoclonal antibodies were found to be dose-dependent. 4C2 was found to have a maximum inhibition of 50%, while the maximum inhibition by F49b was ∼40%. In contrast, two anti-C3c monoclonal antibodies (F39b and F20b) were able to enhance the formation of the factor B-C3b complex in a dose-dependent manner. Competition binding studies with isolated C3 fragments further supported the involvement of the C3d region in factor B binding, as C3d inhibited the binding of factor B to C3b completely. Furthermore, additional competition binding studies conducted with each of the three domains of factor B (Ba, vWF and SP) demonstrated that each domain independently inhibited the binding of intact factor B to C3b by ∼50%.

## Introduction

The complement system plays a major role in innate immunity and consists of a large number of soluble (plasma) and membrane-bound proteins. Depending on the activation trigger, the complement system can be activated via three major pathways, namely, the classical pathway, the lectin pathway, and the alternative pathway. A key characteristic of the alternative pathway is the spontaneous deposition of C3b on the surface of a pathogen, independent of a specific target. During complement activation, C3, the major opsonin of complement, is cleaved to form C3b. Complement factor B, a 90 kD glycoprotein, is a serine proenzyme present in human plasma, and forms a complex with C3b. This renders it susceptible to cleavage by factor D to form two fragments, Ba and Bb. Bb remains transiently bound to C3b forming the C3 cleaving enzyme C3bBb, stabilized by properdin, which activates more C3 [1]. The active site of the C3bBb protease is in the Bb fragment [2]. When examined by electron microscopy, factor B appears to have a three-lobed globular structure [3]. The ∼30 kD, Ba fragment forms the amino-terminal domain of factor B and consists of three CCPs (complement control protein repeats) [4], which are independently-folding modules each approximately 60 amino acids in length [5]. The middle domain of factor B (amino-terminal of Bb) has amino acid sequence homology to a region of about 210 amino acids present in a variety of other proteins which have been named the *A-domain superfamily* [6]). Sadler et al. [7] first recognized the region as a unique domain in von Willebrand Factor which contains three such regions of sequence similarity. For this reason, the domain in factor B has more frequently been termed the von Willebrand Factor domain as will be the case in this paper. The C-terminus of Bb has amino acid sequence homology to serine proteases including highly conserved regions around the active site and the substrate binding site [2, 8]. Investigations of the interaction of factor B with C3b or C3(H2O) have been carried out in a variety of conditions [9-13]. Research carried out in the 1970s pointed to an absolute requirement for Mg^2+^ in the formation of the factor B-C3b complex [14, 15]. Later, Pryzdial and lsenman (1986) [12] demonstrated the formation of a fluid phase C3b-factor B complex in the presence of excess EDTA.

The binding of factor B to C3b involves multiple binding sites within both molecules. The use of synthetic C3 peptides and monoclonal antibodies to C3 has facilitated the mapping of binding sites within the C3 molecule. It is clear that the region at the amino-terminus of the C3b a’ chain is an important region in terms of binding to other complement proteins including factor B (i.e. residues 727-789; within the C3c fragment) [16, 17]. A region spanning residues 933-942 (in the C3g region) has also been implicated as being important [18] in binding factor B. Other studies have been undertaken where monoclonal antibodies to both the C3c and C3d regions could inhibit binding of factor B to C3b [19, 20]. It appears likely that more than one site in C3b interacts (weakly) with factor B such that C3b-B interaction arises from multiple low affinity interactions. The weak interaction or loose arrangement, was also identified by crystal structure analysis of C3bBb, stabilized by the bacterial immune-evasion protein SCIN, where Bb appeared to ‘dangle’ from the tip of the C3b structure [21].

A two-site attachment model has been proposed for the binding of factor B to C3b by several authors [21-24]. One site of contact was depicted as being present in the Ba domain and the other in the Bb domain. Evidence for the existence of a binding site for C3b in the Ba domain has been proposed [22, 23]. Direct evidence for the existence of a binding site for C3b within the serine protease domain has also been reported [25, 26]. Evidence for a role for the von Willebrand Factor domain in binding C3b has also been suggested [3, 21, 26]. Crystal structure analysis of C3b-FH CCP1-4 and C3bBb suggests overlapping binding sites for the CCP-1-2 domains of Factor B and the CCP1-2 domains of Factor H, which leads to displacement of Factor B by steric hindrance. For Factor H, even three binding sides were identified on C3b [27]

Traditional methods for the study of binding of complement components to C3b have involved the use of zymosan or erythrocytes as the solid phase to which C3b becomes attached. Here, we use thiol-Sepharose as a solid phase to overcome the disadvantages of previous methods as thiol-Sepharose is chemically more homogeneous than zymosan or erythrocytes and C3b becomes distributed evenly on its surface which is otherwise not the case [9, 28]. Edens et al. (1990) [29]) studied the effect of several commercially available, pre-activated affinity chromatography supports for their ability to immobilize C3b that would retain functional activity. Functional activity was demonstrated by the conversion of factor B to Bb and Ba in the presence of factor D. Although C3b is bound to this resin via the SH group of the thiolester and not via the carbonyl group, C3b immobilized on thiopropyl activated agarose was concluded to be a suitable support to study C3b interactions. C3b bound to this type of support retained activity compared to fluid phase C3b in terms of factor B consumption. Surface plasmon resonance analysis has detected binding of factor B to C3b and its fragment C3d, but not to the C3c fragment, possibly due to a conformational change in the N-terminal part of C3b during cleavage [30].

In this study, employing C3b-thiol-Sepharose, several parameters of factor B binding to C3b have been investigated including the effect of salt concentration, pH, metal ion concentration and time. The effects of several monoclonal antibodies to the C3c and C3d regions of the C3 molecule on the binding of intact factor B to C3b were also investigated as well as the effects of the individual, purified domains of factor B.

## MATERIALS AND METHODS

### Reagents

Fresh-frozen plasma used for purification of complement components was obtained from the Oxford Regional Blood Transfusion Service, John Radcliffe Hospital, Oxford. The serine protease inhibitor Pefabloc-SC (4-(2-aminoethyl)-benzene-sulphonyl-fluoride) was obtained from Pentapharm AG, Basel, Switzerland. Thiol Sepharose 4B was obtained from Pharmacia, Milton Keynes, Bucks, UK. Lactoperoxidase (L-8257) used for radiolabelling of factor B came from Sigma Chemical Co., Poole, Dorset, U.K as did porcine pancreatic elastase (E-0127) for digestion of factor B to produce the serine protease domain.

### Purification of complement components

Factor B was purified using dye-ligand affinity chromatography, as described by Williams and Sim (1993) [36]. C3 was purified and converted to C3b using a scaled-up version of the method of Dodds (1993) [31]. The final chromatography step of the procedure was increased in scale by replacement of the Mono S 5/5 column with a 1.6 cm x 10 cm diameter pre-packed Pharmacia Hiload Q-Sepharose column which has a higher protein binding capacity. Factor H was purified, as described by Sim et al [32]. C3 fragments, C3c and C3d, were made, as described earlier [33], and factor D was purified as described earlier [34].

### Generation and purification of factor B fragments/domains

The Ba and Bb fragments of factor B were produced by the method described previously [35]. The middle domain of factor B (i.e. the van Willebrand Factor domain, residues 229-444) was produced in high yield using an *E. coli* fusion protein expression system as described Williams and Sim (1994) [36].

The serine protease domain was generated and purified using a modified procedure [25]. Purified factor B (7 mg; 140 µg/ml) in 20 mM tris-HCI, 140 mM NaCl, pH 7.4 was incubated with pancreatic elastase (2% w/w) for 12-16h at 37°C in a final volume of 35 ml. Digestion led to the appearance of a 32 kD fragment, as assessed by a 10% (w/v) SDS-PAGE. N-terminal sequence analysis identified the fragment as the serine protease domain (N-terminal sequence MIDESQSLS). After the overnight incubation, the elastase was inhibited using the serine protease inhibitor Pefabloc-SC (added to 1 mM) and the sample was diluted 10-fold to reduce ionic strength. The serine protease domain was then separated from other minor fragments using ion-exchange on a Mono-S (Pharmacia) FPLC column (1 ml volume) pre-equilibrated in 25 mM Tris-HCI, pH 7.4 at room temperature. After loading the sample, the column was passed with a linear NaCl gradient (total volume 36 ml) to high salt buffer (25 mM Tris-HCI, 500 mM NaCl, pH 7.4) using a flow rate of 1 ml/min. The serine protease domain eluted as a single peak from the Mono-S column at ∼0.3 M NaCl. Pooled, purified fractions were dialysed against PBS and stored in 0.5 ml aliquots at -20°C.

### Radioiodination of factor B

Factor B was radiolabelled by lactoperoxidase-catalysed iodination using the method described earlier [37]. Purified factor B (100 µg in 0.45 ml of PBS) was incubated with 1 µI of lactoperoxidase (1 mg/ml in PBS) and 0.5 mCi (5 µI) of Na^125^I (carrier-free, #IMS 30, Amersham). The reaction was initiated by the addition of 1 µI of 0.03% H_2_O_2_ (30% H_2_O_2_ stock solution diluted x 1000 in distilled water). The mixture was then shaken gently and incubated for 30 mins on ice. ^125^I-labelled protein was desalted on a PD10 column (Pharmacia) pre-equilibrated with 10 mM Pipes [(Piperazine-N, N’-bis-2-ethanesulphonic acid) (sodium salt), pH7.0. Using this method, factor B was radiolabelled to a specific activity of 1 x 10^6^ cpm/µg.

### Preparation of C3b-thiol-sepharose

The assay system developed to study the various parameters of factor B binding to C3b involved the binding of C3b to thiol-Sepharose via the free SH group in C3b. C3 was purified and converted to C3b, as reported [31]. The sample was reacted with methylamine so that any remaining intact C3 would be converted to C3(CH_3_NH_2_) [38], as follows: material was dialysed against 0.5 M Tris-HCI, pH8.5 overnight at 4°C. Methylamine (solid) was added to the sample to 25 mM and incubated for 1h at 37°C. Subsequent dialysis against 25 mM Tris-HCI, 140 mM NaCl, pH 8.2 was carried out to remove the methylamine in preparation for binding to thiol-Sepharose. Approximately 4 g of dry thiol-Sepharose 48 resin was pre-swollen in 40 ml of PBS for 2h at room temperature following the manufacturer’s instructions. 1 g swells to approximately 4 ml. The resin was then washed with 200 ml of PBS on a sintered glass funnel before reduction with 50 mM OTT at 37°C for 30 min. After a further washing step, the resin was resuspended as a 1:1 (v/v) slurry in 25 mM Tris-HCI, 140 mM NaCl, pH8.2. C3b (0.5 mg per ml of packed resin) was added to the thiol-Sepharose resin and incubated at room temperature for 2h using a rotary stirrer. Using the above concentrations of C3b, approximately 20-25% of the protein bound to the resin. Thus, the content of C3b was routinely 100-125 µg/ml of resin. The percentage of protein bound was estimated by measuring the OD_280_ of the original solution and of three subsequent washes with PBS. After the washing steps, any free SH groups on the resin were blocked by incubation of the resin with 5 mg of iodoacetamide per ml of resin for 1h using a rotary stirrer. Non-specific binding sites on the C3b-thiol-Sepharose were blocked by incubation of the resin with 1 mg of BSA per ml of resin for 1h at room temperature. Control resin was subjected to the same treatment, but without addition of C3b.

### Direct binding of ^125^I-labelled factor B to C3b-thiol-Sepharose

The C3b-bearing and control resins were used as a 1:1 (v/v) slurry (i.e. 1 volume of packed resin: 1 volume of buffer) in binding buffer [10mM Pipes (sodium salt), 1mg/ml BSA, pH 7.0 containing either 0.2 mM MgCl_2_ or 5 mM EDTA]. For each binding reaction, 100 µI of slurry was added to a 1.5ml Eppendorf tube. ^125^I labelled factor B was added to each tube (2-5 x10^5^ cpm) and the tubes were incubated at room temperature (generally for 1 h). All experiments were carried out in duplicate. After incubation, the supernatant was aspirated off and 200 µI of a 1:1 (v/v) slurry of underivatised Sepharose 4B in binding buffer was added to help prevent loss of C3b-thiol Sepharose resin during washing steps. The resin was washed 5 times with 0.85 ml of binding buffer by centrifugation at 2500 rpm for 3 min after each wash. After the final washing step, the resin was resuspended in 150 µI of binding buffer and 200 µI of the resin/buffer suspension was counted in an LKB 1275 Mini-gamma counter (counting efficiency∼ 68%). Direct binding of ^125^I-labelled factor B to C3b-Sepharose was investigated in a range of pH values, salt concentrations, magnesium ion or EDTA concentrations and for varying lengths of time. In each case, washing steps were carried out in the appropriate bindingbuffer. Direct binding of individual domains of factor B radiolabelled using the lactoperoxidase-catalysed method of iodination was also tested using the method above.

### Assays to study the effects of monoclonal antibodies against C3 on the binding of ^125^I-labelled factor B to C3b-thiol Sepharose

A number of monoclonal antibodies raised against the C3c and C3d regions of C3 [19, 39] were purified from ascites fluid by triple 18% (w/v) sodium sulphate precipitation [40]. The purified antibodies were dialysed against and used in PBS buffer. Preliminary assays were carried out to investigate the effect of each monoclonal antibody on formation of the factor B-C3b complex. The resin was used as a 1:1 (v/v) slurry in PBS. For each monoclonal antibody tested, 100 µI slurry was incubated with 50 µg of the appropriate purified monoclonal antibody in 150 µI of PBS. The tubes were incubated overnight at 4°C and the resin was then washed 5 times with 0.85 ml of PBS. After the final wash, the C3b-thiol Sepharose (50 µI) was resuspended in 50 µI of 10 mM Pipes (sodium salt), 0.2 mM MgCl_2_, 1 mg/ml BSA, pH 7.0, and direct binding of ^125^I-labelled factor B was assessed, as described above. The anti-factor H monoclonal OX23 [41] was used as a negative control. A rabbit anti-C3d polyclonal raised by immunisation with C3d, purified by the method of Micklem et al. [33], was used as a positive control. It was used in the form of lgG purified by triple 16% (w/v) sodium sulphate precipitation. Four monoclonal antibodies were found to have an effect and were selected for further study. Two monoclonal antibodies to the C3c region of the C3 molecule (F20b and F39b) were selected. These antibodies were found to increase the amount of factor B bound to the C3b-thiol Sepharose compared to samples where pre-incubation of C3b-thiol Sepharose had been carried out in the presence of the OX23 negative control monoclonal antibody, or in the absence of monoclonal antibody. In contrast, pre-incubation of the C3b-thiol Sepharose with two monoclonal antibodies to the C3d region (F49b and 4C2) resulted in inhibition of the binding of factor B to C3b-thiol Sepharose. The dose-dependent effect of these monoclonal antibodies was assessed as described below.

In each case 100 µI of C3b-thiol Sepharose slurry in binding buffer was pre-incubated with serial 2-fold dilutions of monoclonal antibody (200 µI; max. final concentration 670 µg/ml in PBS) overnight at 4°C. After the overnight incubation of C3b-thiol-Sepharose resin with antibody the resin was washed 5 times with 0.85 ml of PBS. Binding of 125I-labelled factor B was assessed as described above. Binding was tested in the presence of control antibodies as well as in the absence of antibody.

### Assay to examine the effects of potential competitors (factor H, factor B domains/fragments and C3 fragments C3c and C3d) on the binding of factor B to C3b

In each assay 100 µI of C3b-thiol Sepharose 1:1 (v/v) slurry in binding buffer was pre-incubated with serial two-fold dilutions. of potential competitors of C3b-factor B binding: these included factor H (400 µI; max. concentration 125 µg/ml), Ba, Bb, the serine protease domain or vWF domain (200 µI; max. concentration 125 µg/ml). A control with no potential competitor was included as well as controls with an irrelevant protein (ovalbumin) as potential competitor^. 125^I-labelled factor B (4 x 10^5^ cpm) was then added and the radioactivity bound measured after a 1h incubation at room temperature and five washes with binding buffer as above.

Other competition assays were set up where a constant quantity of ^125^I-labelled factor B (7 µI; concentration 50 µg/ml) was pre-incubated with serial 2-fold dilutions of C3b, C3c or C3d (100 µI; max. conc. 100 µg/ml) for 2 h at room temperature in 10mM Pipes (sodium salt) buffer, pH 7.0 containing 0.2 mM MgCl2 and 1mg/ml BSA. Parallel assays were set up with C3b as a positive control and ovalbumin or no competitor as a negative control. C3b-Sepharose (100 µI of slurry in binding buffer) was then added to each tube. After incubation and washing (as above), the ^125^I-labelled factor B bound in the presence and absence of competitor was measured.

### Examination of the effect of factor D on the C3b-factor B interaction

The effect of factor D on the factor B-C3b complex was investigated by the following method. ^125^I-labelled factor B (50 µl; 50 µg/ml) was incubated with 200 µI of C3b-thiol Sepharose suspended in 200 µI of binding buffer containing 0.2 mM Mg^2+^ or 5 mM EDTA for 1 h at room temperature. This was followed by five washes as above and resuspension of the resins in 200 µI of binding buffer. At this point, 1.5 µg of factor D (3 µI of a purified factor D solution at a concentration of 0.5 mg/ml in binding buffer) was added to the resin/buffer suspension in each tube. The tubes were then incubated at 37°C with the removal of aliquots of resin/buffer suspensions at appropriate time points (5, 25 and 50 mins). Controls were included which contained ^125^I-labelled factor B bound to the C3ba thiol Sepharose without the addition of factor D. A further: control was ^125^I-labelled factor B with the addition of factor D in the absence of C3b-thiol Sepharose. Aliquots (100 µl of resin/buffer suspension) were removed at appropriate time points and the supernatants added to SDS-PAGE sample buffer. The samples were analysed by SDS-PAGE on 7.5% (w/v) polyacrylamide gels to assess breakdown of factor B to Bb and Ba fragments. The gels were dried and autoradiographed overnight. The bands observed on the developed autoradiographs were scanned using a Molecular Dynamics computing densitometer.

## RESULTS

### Direct binding of 125I-labelled factor B to C3b-thiol Sepharose

Radioactivity bound to control resin was always low compared to that bound to C3b-thiol Sepharose (∼0.1%). For this reason, background counts were always subtracted from the binding results shown. ln **Fig. 1a**, it is shown that factor B binds to C3b-thiol Sepharose in a saturable manner. Assuming that factor B and C3b interact in a 1:1 ratio then it can be calculated from the graph that 0.1% of the C3b present is available for binding. This figure appears low, but is not unusual in experiments with porous matrices: immobilized trypsin, in a similar system, for example, has only about 0.5% activity relative to soluble trypsin in cleaving protein substrates [28].

**Fig. 1.**
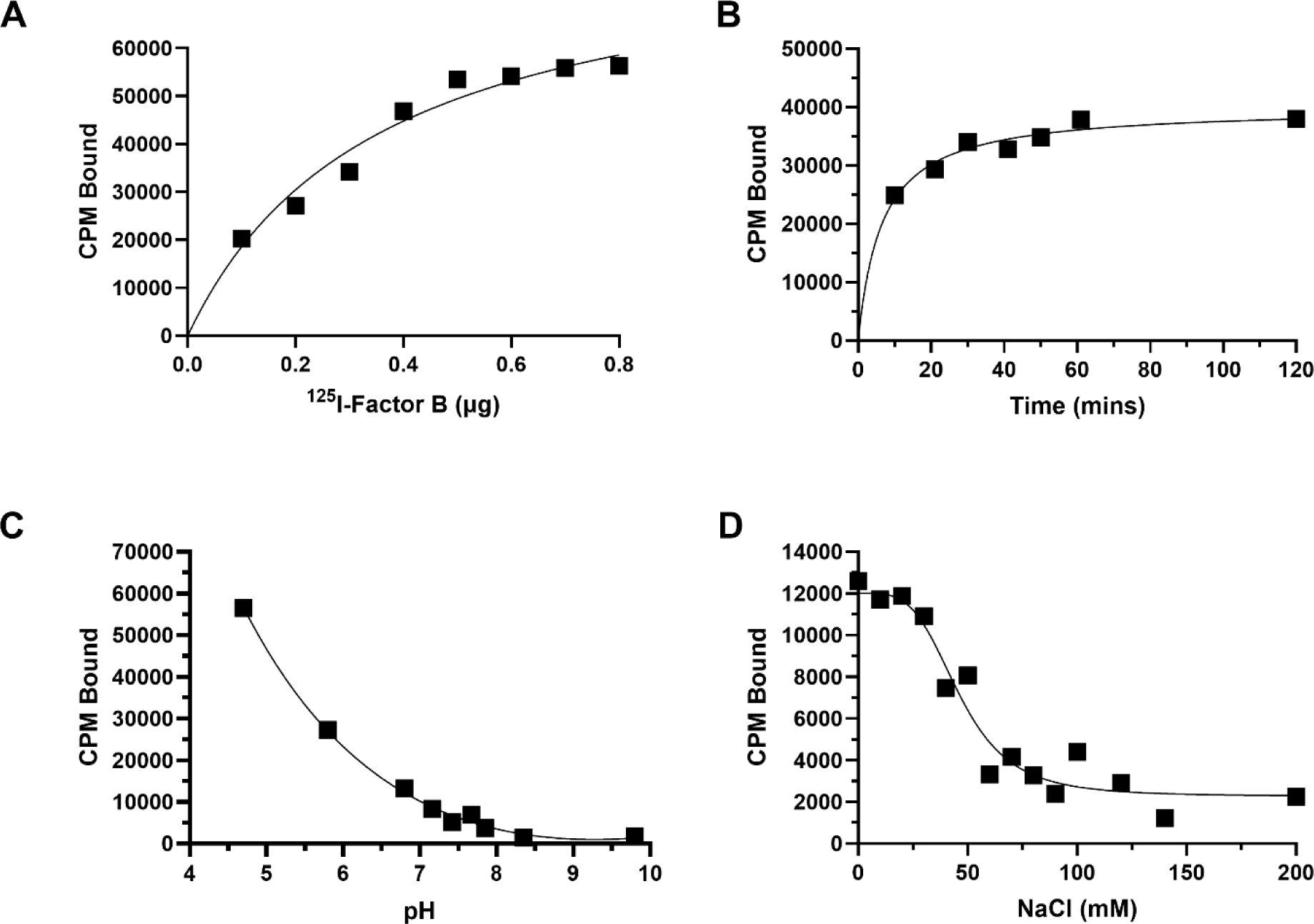
Binding of 125I-labelled factor B to C3b-thiol Sepharose. a) Demonstration of saturation: the assay was carried out as described in the methods section except that increasing amounts of ^125^I-labelled factor B (0.2-1 µg; 0.2-1 x 106 cpm) were incubated with 25 µI of C3b-thiol Sepharose suspended in 25 µI of binding buffer. b) Time-dependence of binding: the assay was carried out as described in the methods section except that tubes were incubated for varying lengths of time. c) pH-dependence of binding: a number of buffers of pre-adjusted pH values made up of 20 mM MES, 20 mM HEPES, 20 mM tris and 20 mM glycine (pH adjusted with NaOH or HCI) were diluted x 4 with distilled water to reduce the total buffer concentration to 20 mM. The pH values were as follows: 4.7, 5.8, 6.8, 7.16, 7.42, 7.67, 7.85, 8.35 and 9.8. d) Effect of ionic strength on binding: the assay was carried out as described in the methods section except that incubation was carried out in binding buffer containing different concentrations of NaCl.

**Fig. 1b** shows the time-dependence of binding of ^125^I-labelled factor B to C3b-thiol Sepharose. The amount of ^125^I-labelled factor B bound to C3b-thiol Sepharose did not increase after a period of 60 min at room temperature. The relatively long time period required for maximum complex formation to occur may reflect the porous nature of the C3b-thiol Sepharose such that the time required for the factor B molecules to diffuse to all accessible C3b molecules was relatively long.

In **Fig. 1c**, the effect of pH on the binding of ^125^I-labelled factor B to C3b-thiol Sepharose resin is shown. The highest binding appears to take place at the lowest pH value i.e. pH 4.7 and that binding decreased rapidly from pH 4.7-6.8. Assays were also carried out to study the effect of different concentrations of NaCl on the factor B-C3b interaction in this system (**Fig. 1d).** Maximum binding took place at low ionic strength. The binding did not decrease dramatically from 0-30 mM NaCl, but from 30-60 mM NaCl, the binding does decrease significantly.

The binding of ^125^I-labelled factor B to C3b-thiol Sepharose in various Mg^2+^ and EDTA concentrations (0-5 mM) was also measured and the results are shown in **Fig. 2**. Binding of factor B took place in the presence of Mg^2+^, and also to a lesser extent, in the presence of EDTA. In 5 mM Mg^2+^, the binding was greater by 50% than the binding in 0.02 mM Mg^2+^. Binding in the presence of EDTA remained approx. the same over the 0.02-5 mM EDTA concentration range and was always lower than in Mg^2+^. In 5 mM EDTA, binding of ^125^I-labelled factor B to C3b-thiol Sepharose was always observed to be approx. 60-70% of that in 0.2 mM Mg^2+^.

**Fig. 2.**
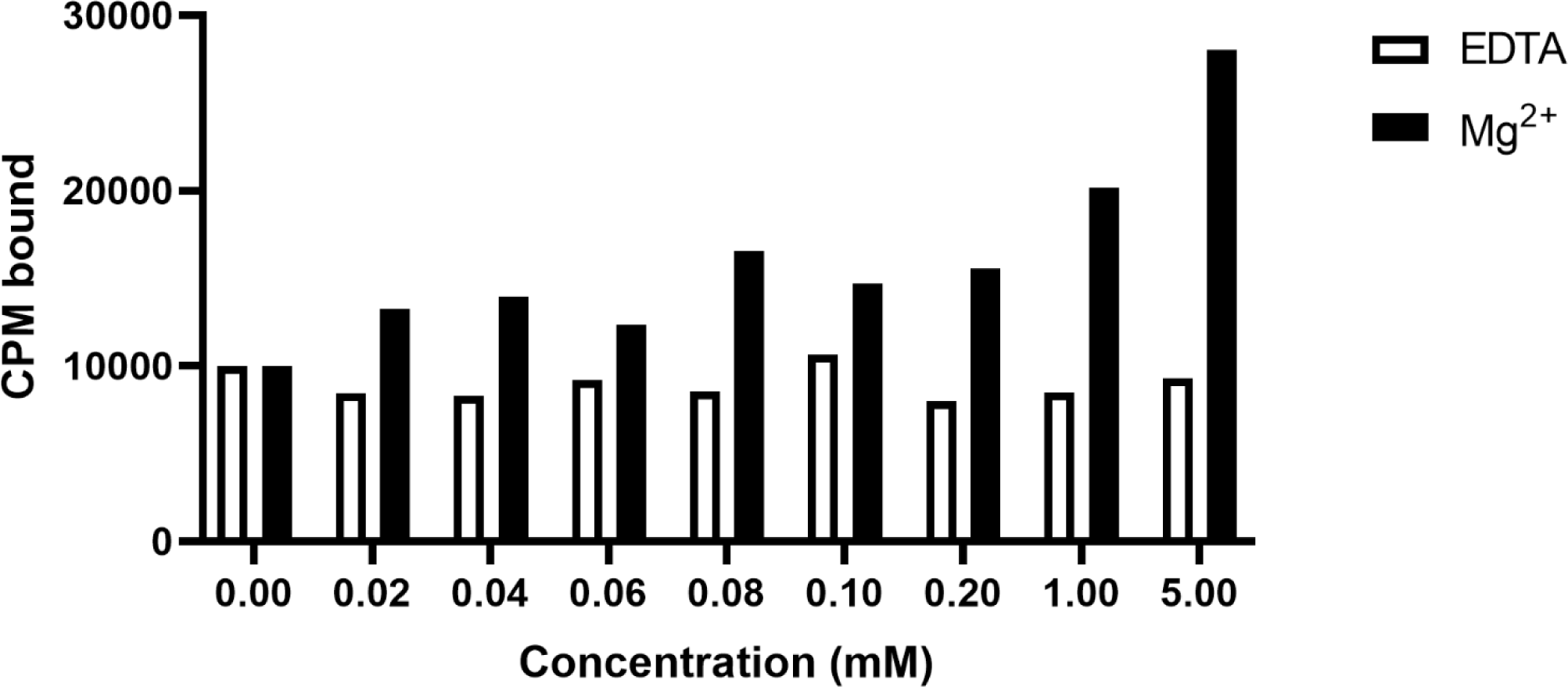
Investigation of the effect of Mg^2+^ and EDTA concentration on the C3b-factor B complex. The assay was carried out as in the methods section except that binding was carried out in the following concentrations of either MgCl_2_ (▪) or EDTA (□): no added Mg^2+^ or EDTA, 0.02, 0.04, 0.06, 0.08, 0.1, 0.2, 1 and 5 mM.

### The effect of anti-C3 monoclonal antibodies on the factor B-C3b interaction

Assays to study the effect of several monoclonal antibodies to C3 on the factor B-C3b interaction were carried out, as described in the methods section. The purity of the antibodies selected for further study as well as control antibodies is shown in **Fig. 3**, as assessed by SDS-PAGE on a 7.5% (w/v) polyacrylamide gel. Maximum observed inhibition (F49b and 4C2) or enhancement (F20b and F39b) of factor B binding to C3b-thiol Sepharose is shown in **Table 1**. Observed effects were dose-dependent (data not shown). C3b-thiol-Sepharose resin was also incubated with the rabbit anti-C3d polyclonal lgG fraction (positive control). The maximum inhibition of factor B binding to C3b incubated with this polyclonal positive control was ∼80% (at an approximate molar excess of specific anti-C3d antibodies of ∼50 fold assuming that 2% of the lgG preparation was C3d-specific). The failure to achieve complete inhibition may lie in the inability of multiple antibodies to bind to each C 3 b molecule due to spatial constraints imposed by the Sepharose matrix.

**Fig. 3.**
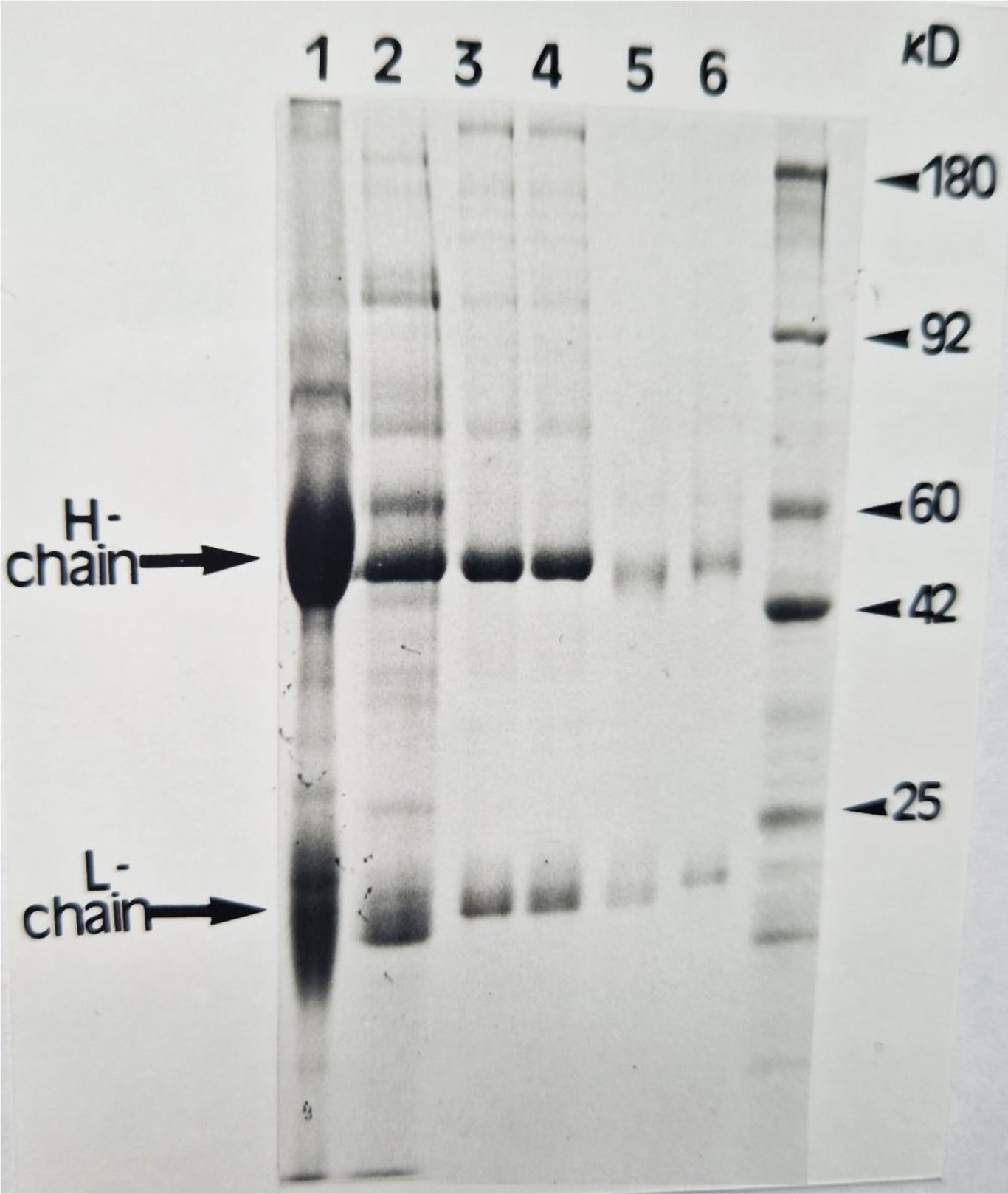
SDS-PAGE analysis (7.5%w/v gel) of purified antibodies to C3. Monoclonal antibodies were purified by triple 18% (w/v) sodium sulphate precipitation and polyclonal antibodies to C3d were purified by triple 16% (w/v) sodium sulphate precipitation. All samples were dialysed against PBS at 4°C overnight. Aliquots of each antibody solution were run under reducing conditions and proteins were visualized by staining with Coomassie Brilliant Blue. The identities of the tracks are as follows: lane 1-anti-C3d polyclonal; lane 2-4C2; lane 3-F49b; lane 4-F39b; lane 5-F20b and lane 6-OX23.

**Table 1.** Effect of anti-C3 antibodies on the interaction of factor B with C3b-thiol Sepharose. Results shown represent the effects of antibodies at maximum concentrations of 200 µg/ml and in the presence of 0.2 mM Mg^2+^

| **Monoclonal Antibody** | **Inhibition or enhancement of factor B binding to C3b-thiol Sepharose** |
| --- | --- |
| 4C2 | 52% Inhibition |
| F49b | 44% Inhibition |
| F20b | Enhancement of binding, x1.7 as much factor B bound as compared with pre- incubation of C3b-thiol Sepharose with OX23 control monoclonal antibody. |
| F39b | Enhancement of binding as above except x2.8 as much factor B bound. |
| anti-C3d polyclonal (+ve control) | 80% Inhibition |
| OX23 (-ve control) | No effect |

Maximum inhibition of factor B binding by 4C2 was found to be ∼50% and inhibition by F49b was found to be ∼40% (at an approximate 50-fold molar excess of antibody over C3b). 4C2 is an antibody previously characterized by Koistinen et al. [19], who found that it inhibited the binding of ^125^I-labelled factor B to C3b bound to erythrocytes by up to 80%. The lower percentage of maximum inhibition observed in this study may be explained by the inability of the antibody molecule, diffusing through the porous matrix, to reach as many C3b molecules as the smaller factor B molecule. The anti-C3c monoclonal antibodies enhanced the binding of ^125^I-labelled factor B to C3b-thiol Sepharose. After pre-incubation of 50 µI of C3b-thiol Sepharose (∼ 5 µg of C3b) with 200 µg of F39b (an approximate 50 times molar excess), the amount of ^125^I -labelled factor B which bound was found to be almost three times (x 2.8) as much as when pre-incubation had been carried out with the negative control, OX23 or in the absence of monoclonal. Pre-incubation of 50 µI of C3b-thiol Sepharose with the same quantity of the F20b monoclonal resulted in almost twice (x 1.7) as much factor B binding compared with pre-incubation with the same quantity of 0X23 or in the absence of monoclonal. The effects of the antibodies noted above were in assays with 0.2 mM MgCl_2_ present.

Assays were done to investigate. whether the presence of EDTA changed the effects of the monoclonal antibodies. It was shown that the enhancing effect of the anti-C3c monoclonals (F39b and F20b) was observed in binding conditions containing EDTA as well as in the presence of Mg^2+^. However, the inhibitory effects of the anti-C3d monoclonals (4C2 and F49b) were not observed when binding of factor B was assessed in the presence of 5 mM EDTA (not shown). A possible explanation for this observation could be that EDTA had an effect on the binding of these monoclonal antibodies to the epitopes which they bind in C3d. It was also investigated whether pre-incubation of the C3b-thiol Sepharose with a combination of the two anti-C3d monoclonal antibodies, or the two anti-C3c monoclonal antibodies, would result in additive effects, but no additive effect was seen. The observed effects may be due to the inability of two antibodies to bind to a single C3b molecule due to the porous nature of the Sepharose resin, or possibly that the pairs of antibodies recognized the same or overlapping epitopes.

### Effects of factor H and C3 fragments on the binding of factor B to C3b-thiol Sepharose

Analysis of the purity of the components used in the assays is shown in **Fig. 4a** on a 7.5% (w/v) polyacrylamide gel. The C3d contained traces of C3c. The assay used to study the effect of factor H on the factor B-C3b interaction in the presence of 0.2 mM Mg^2+^ is described in the methods section and the results obtained are shown in **Fig. 4b**. At an approximate 12 times molar excess of factor H over C3b, inhibition of 125l-labelled factor B binding was ∼50%: inhibition had not reached a maximum as a plateau was not observed. It is possible that a larger degree of inhibition could be achieved with higher concentrations of factor H. This result was obtained in the presence of 0.2 mM Mg^2+^ and similar results were obtained in the presence of 5 mM EDTA.

**Fig. 4.**
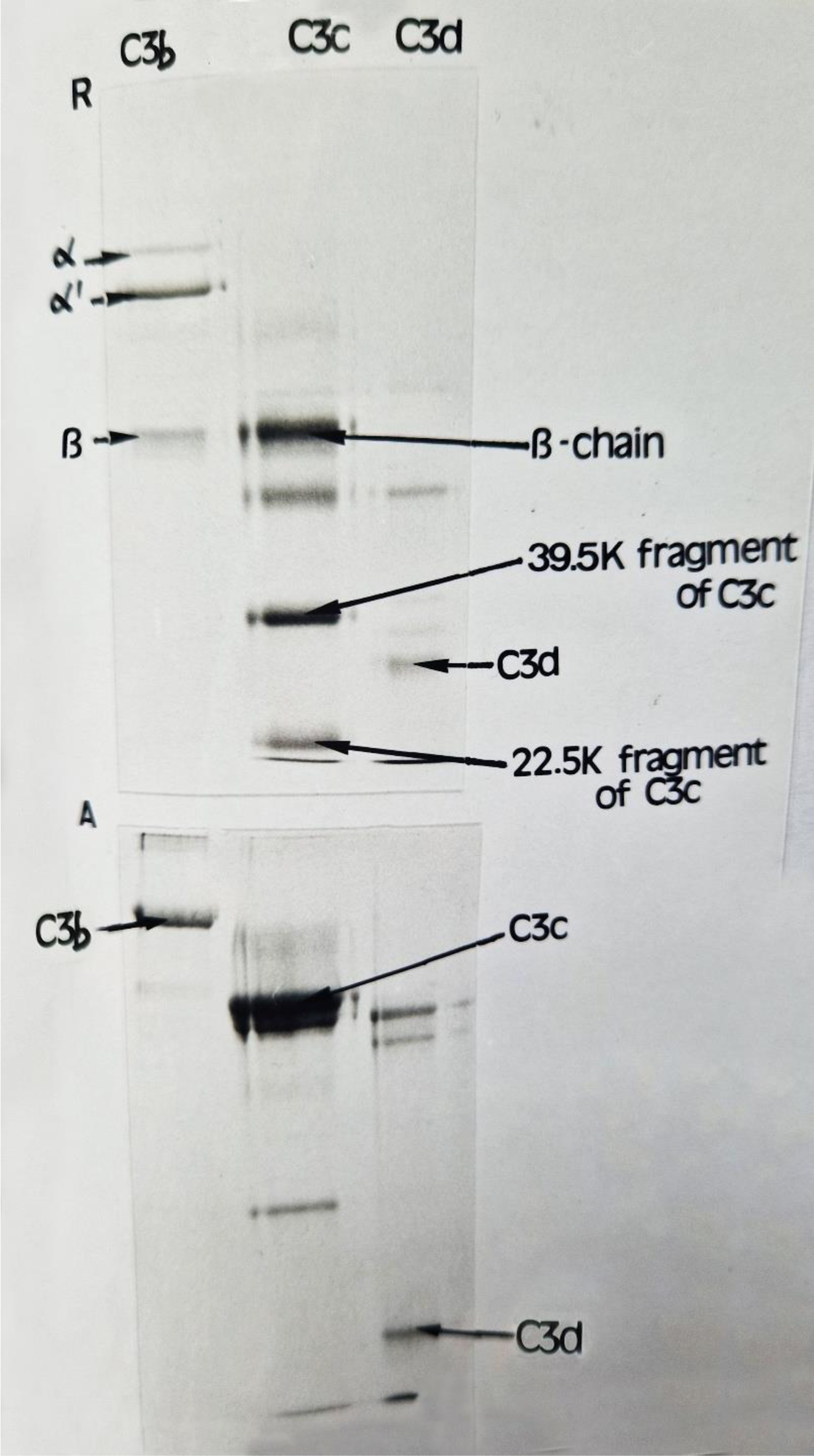
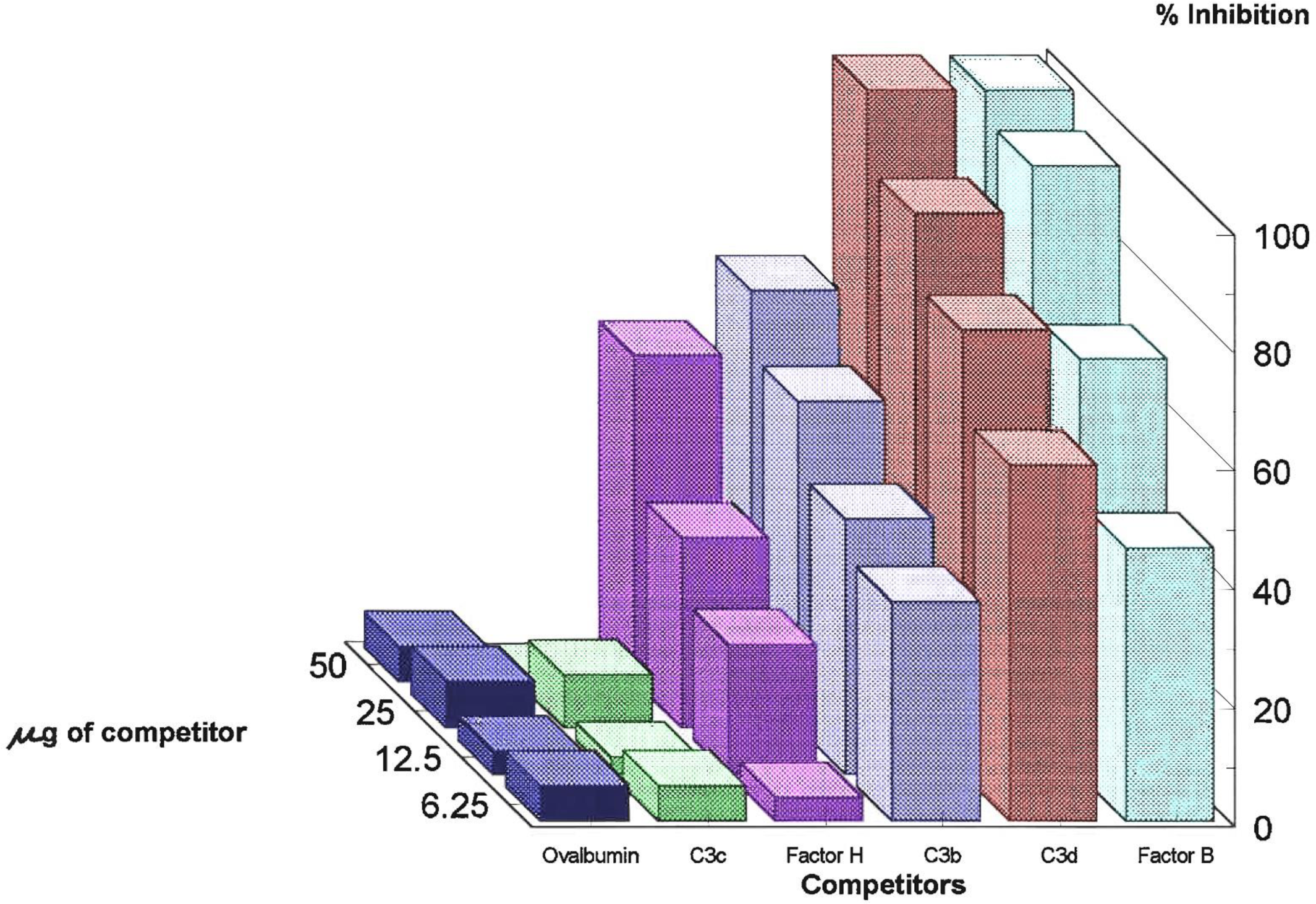
a) **SDS-PAGE analysis (7.5% gel) of C3 fragments** C3 fragments were run reduced (upper panel-A) and non-reduced (lower panel-A) and proteins were visualized by Coomassie Brilliant Blue staining. Cross-contamination of C3d with C3c is evident. b) **Competition binding study of the effect of factor H and C3 fragments on the binding of factor B to C3b-thiol Sepharose** Effects of ovalbumin, C3c, factor H, C3b, C3d and factor B on the binding of factor B to C3b are shown. Tests were done as described in the methods section.

The effects of C3b, C3c and C3d were investigated as described in the methods section. The results obtained in the presence of 0.2 mM Mg^2+^ are also shown in **Fig. 4b**. Pre-incubation of C3b at a 75-fold molar excess over ^125^I-labelled factor B prior to incubation with C3b-thiol Sepharose resulted in 70% inhibition of binding. This amount of fluid phase C3b (50 µg) also corresponded to a 10-fold molar excess over the C3b bound to thiol Sepharose. From **Fig. 4b** it is clear that inhibition of factor B binding to C3b-thiol Sepharose by fluid phase C3b had not reached a maximum.

Pre-incubation of ^125^I-labelled factor B with the same quantity of C3d (50 µg) resulted in almost complete inhibition of binding of factor B to C3b-thiol Sepharose. This quantity of C3d corresponds to a 290-fold molar excess over factor B and a 40-fold molar excess over thiol-Sepharose bound C3b. It was concluded that C3d was the fragment which was exerting an effect and not the contaminating C3c in the preparation as the purified C3c fragment alone had no effect **(Fig. 4b).** Thus, the inhibition observed with C3b appears to be due to the C3d portion of C3b. An assessment of the effects of C3b, C3c and C3d in 5 mM EDTA was also made and similar results were obtained.

### Investigation of binding sites in the Individual domains of factor B for C3b

SDS-PAGE analysis to show the purity of factor B and its fragments/domains on 10% (w/v) polyacrylamide gels is shown in **Fig. 5a**. Direct binding of radiolabelled fragments/domains to C3b-thiol Sepharose was first assessed. Samples of 100 µg of each domain [in 100-500 µI of 10 mM Pipes (sodium salt), 30 mM NaCl, pH7.0] were radio-iodinated using the lactoperoxidase method. Assays of direct binding were carried out as described for the binding of intact ^125^I-labelled factor B to C3b-thiol Sepharose. No direct binding of ^125^I-labelled Ba, Bb, vWF or SP could be demonstrated. Binding of each of the domains/fragments was then studied indirectly by investigating the ability of each to compete with intact ^125^I-labelled factor B for binding to C3b-thiol Sepharose.

**Fig. 5.**
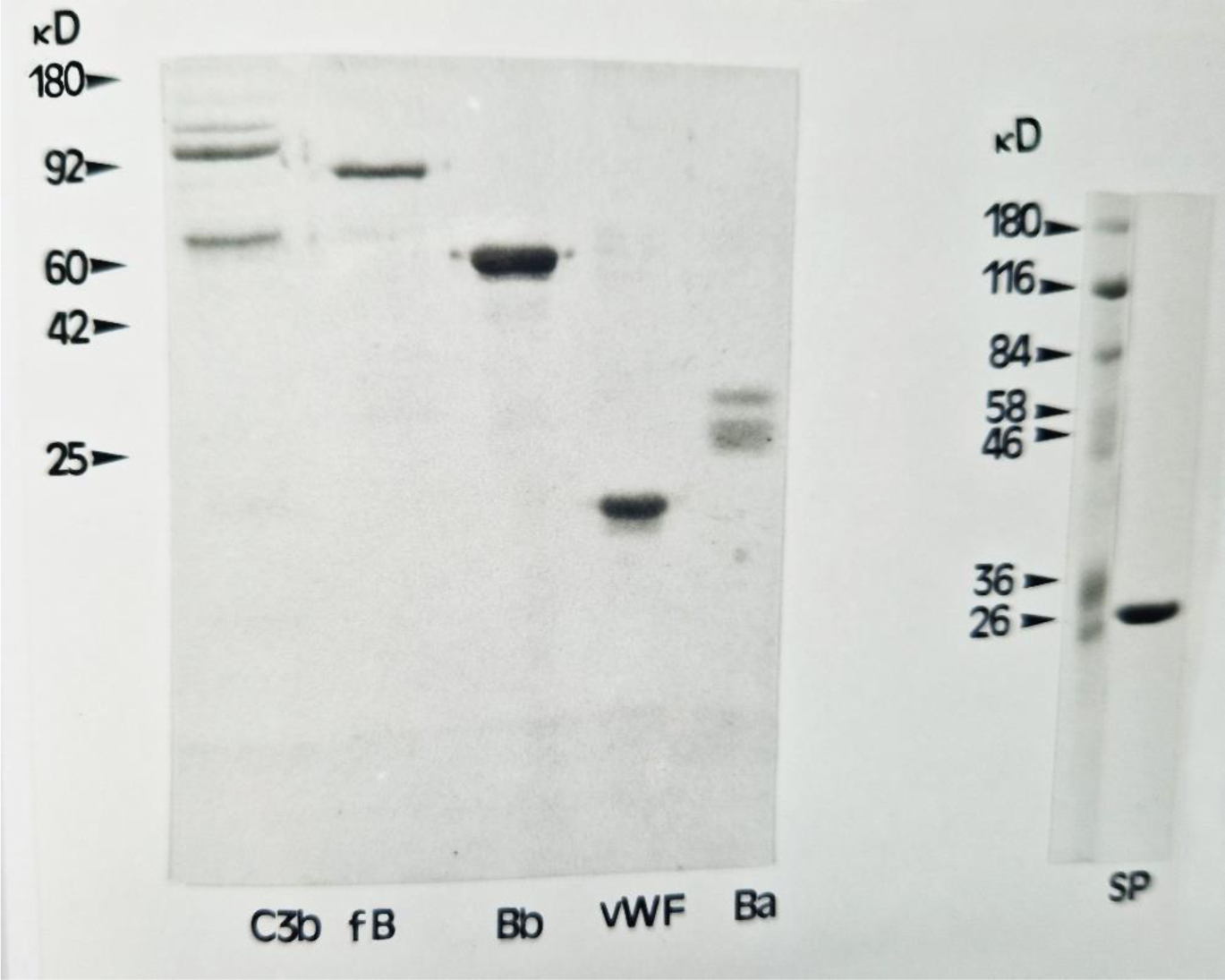
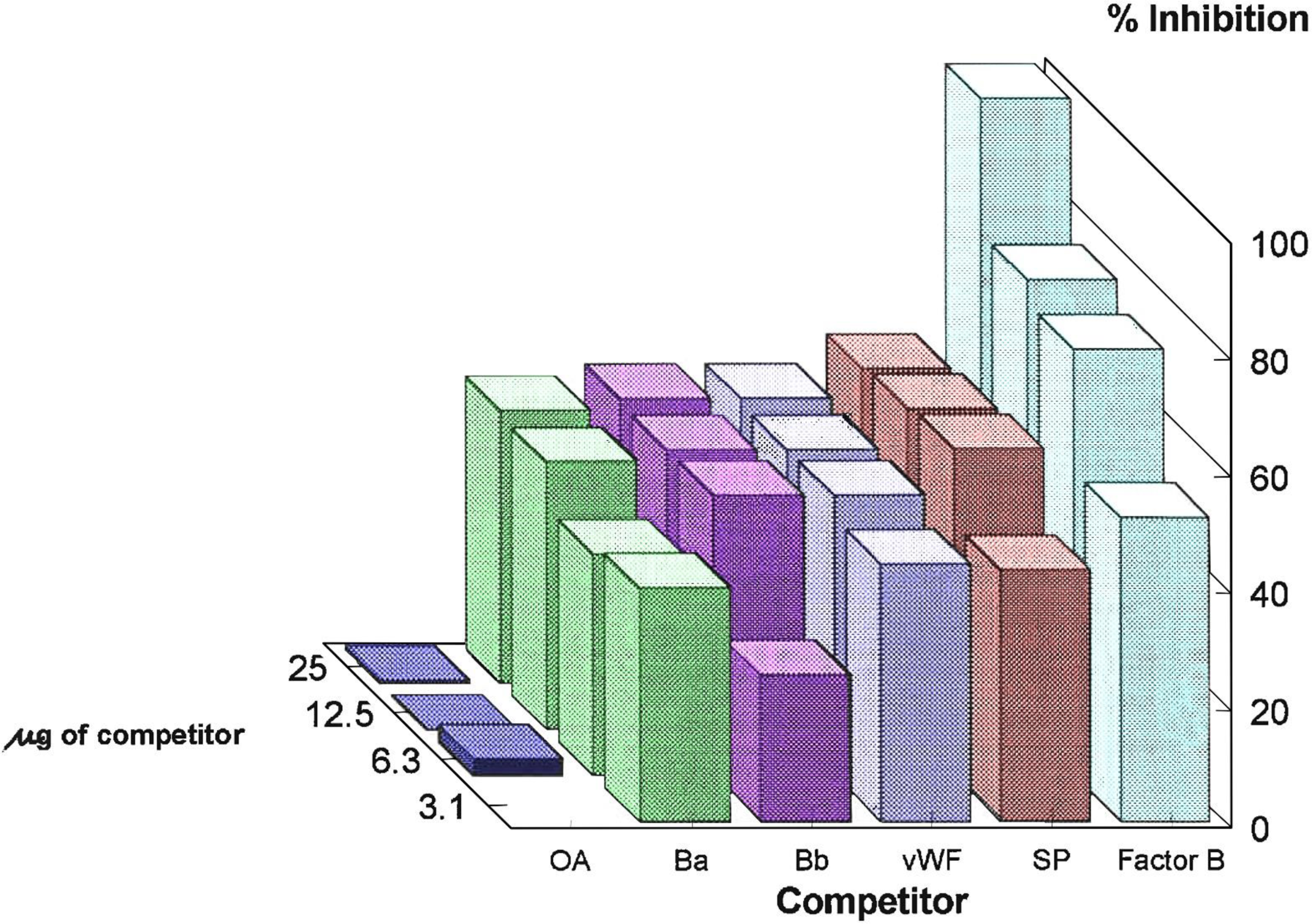
Effects of factor B domains on factor B-C3b interaction. a) Analysis of the purity of Ba, Bb, the serine protease domain (SP), and the von Willebrand Factor domain (vWF) by SDS-PAGE on a 10% (w/v) polyacrylamide gel under reducing conditions. The tracks are identified by the abbreviated terms for the domains or fragments. The gel also shows the purified C3b which was bound to the thiol Sepharose. The proteins were visualized by Coomassie Brilliant Blue staining. b) Competition binding study to investigate indirect binding of the Ba, Bb, vWF and SP fragments of factor B to C3b-thiol Sepharose in the presence of 0.2 mM Mg^2+^.

The Ba and Bb fragments of factor B were able individually to compete with intact factor B for binding to C3b immobilized on thiol Sepharose, both in the presence of 0.2 mM Mg^2+^ **(Fig. 5b)** and in the presence of 5 mM EDTA (not shown). inhibition of ^125^I-labelled factor B binding to C3b thiol Sepharose by intact unlabelled factor B was found to be close to 100% as expected. Ovalbumin (the negative control) had no effect on the binding reaction **(Fig. 5b**). Maximum inhibition by both the Ba and Bb fragments was ∼50%. This was in the presence of 25 µg of Ba or Bb. For Bb, this was an approximately 15-fold molar excess over C3b, while for Ba, it was a 30-fold molar excess. Also, in **Fig. 5b** is shown the effect of the vWF domain as a competitor for binding to C3b. The maximum inhibition achieved at an approximately 30-fold molar excess of vWF over C3b was also ∼50% either in the presence of Mg^2+^ or EDTA. It was also found that the isolated serine protease (SP) domain of factor B was also able to compete with factor B for binding to C3b. The results are also shown in **Fig. 5b**. Up to 50% inhibition was achieved in the presence of magnesium ions and up to 40% in the presence of EDTA with an approximate 30 times molar excess of SP over C3b.

From the results obtained, it appears that each of the three domains of factor B has binding affinity for C3b. The fact that direct binding to C3b was not observed with isolated Ba, Bb, vWF or SP indicates that binding of the individual domains to C3b is weak, and that only when all of the necessary binding sites are present in one ligand molecule can direct binding be observed i.e. in the intact factor B molecule.

The combined effects of a mixture of Ba and Bb was studied by incubating C3b-thiol Sepharose with a sample of factor B (100 µg) which had been digested with trypsin (2% w/w) for 16 min at 37°C. The mixture of fragments was also found partially to inhibit (to ∼50%) the binding of factor B to C3b-thiol Sepharose.

### Effect of factor D on the C3b-factor D interaction

The effect of factor D on pre-formed factor B-C3b-thiol Sepharose complex was studied. The data obtained by scanning of autoradiographs is shown in Fig. 6. After only 5 mins of incubation at 37°C, ∼80% of the radioactivity originally present in factor B bound to the C3b-thiol Sepharose was released into the supernatant as intact factor B (8% of radioactivity), Bb (34%) or Ba (38%) (**Fig**. **6a**). Similar observations were made at 25 and 50 min; thus, the reaction was rapid and essentially completed at 5 min. Similarly, when the experiment was carried out in the presence of EDTA **(Fig 6b),** after only 5 min of incubation of factor B-C3b thiol Sepharose with factor D, approximately 82% of the radioactivity present in factor B which had been bound to the resin was released into the supernatant as intact factor B (12%), Bb (38%) or Ba (32%).

**Fig. 6.**
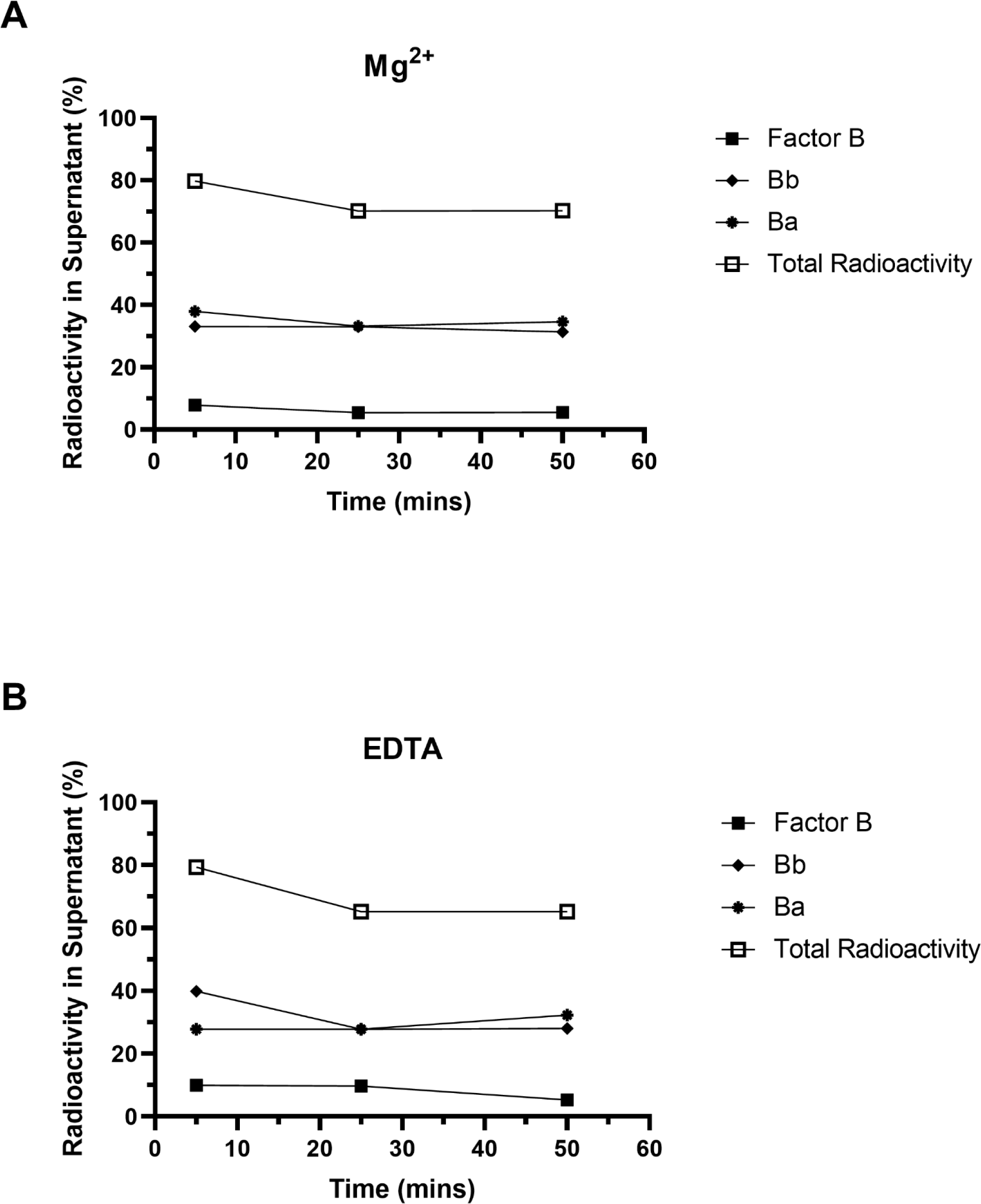
quantitation of factor B. Bb and Ba fragments present in the supernatant after cleavage of factor B bound to C3b-thiol Sepharose. The autoradiographs obtained from the experiments described in the methods section in the presence of Mg^2+^ (a) or EDTA (b) were scanned and the quantity of factor B (▪), Bb(♦) and Ba (*) bands present in the supernatants are expressed as percentages of the quantity of factor B initially bound to the resin. The total radioactivity in the supernatant is represented by (□).

## DISCUSSION

The binding of ^125^I-labelled factor B to C3b-thiol Sepharose provides a new assay system which has been used to study the C3b-factor B complex. Direct saturable specific binding of ^125^I-labelled factor B to C3b thiol Sepharose was demonstrated **(Flg.1a)** and the system was used to investigate various parameters of binding. The effect of pH in the range 4.7-9.8 on the formation of the complex was investigated and it was found that complex formation was more extensive at lower pH values **(Fig. 1c**). Fishelson et al (1987) [42] previously studied the binding of factor B to C3b on red cells in a narrower pH range (6.0-8.0), and a similar effect at lower pH was observed. The effect of increasing ionic strength on the factor B-C3b thiol Sepharose complex is also reported here **(Fig. 1d).** Maximum binding took place at low ionic strength and binding decreased dramatically from 30-60 mM NaCl. DiScipio (1981) [9] studied the effect of ionic strength on the binding of ^125^I-labelled factor B to C3b’ “zymosan. Although the measurements carried out by DiScipio et al were carried out over a wider range of NaCl concentrations (0-600 mM NaCl), the results were consistent with those reported here.

Pryzdial and lsenman (1986) [12] and Koistinen et al. (1989) [19] demonstrated some formation of a fluid phase C3b-factor B complex in the presence of EDTA. These data contradicted earlier research in the 1970s which pointed to an absolute requirement for Mg^2+^ in the formation of the complex [14, 15, 43]. However, some observations could be due to buffer conditions differing from those reported here. With the use of a very sensitive functional assay [12], it was also reported a Mg^2+^-independent interaction for the binding of factor B to solid-phase C3b in excess EDTA. Here, the binding of factor B to solid phase C3b (bound to thiol Sepharose) was readily detectable in the presence of EDTA **(Fig. 2).** Mg^2+^ did appear to have an enhancing role on the binding of factor B to C3b as binding in the presence of Mg^2+^ was always higher (by 30-40%) than the binding observed in the presence of EDTA. These results, however, may cast doubts upon the belief that Mg^2+^ is an absolute requirement for formation of the factor B-C3b complex.

Evidence was also presented that two anti-C3d monoclonal antibodies (F49b and 4C2) were able partially to inhibit the binding of factor B to C3b **(Table 1**). Koistinen et al. (1989) [19]) previously reported that 4C2 was able to inhibit the binding of factor B, factor H and CR1 to C3b. The fact that both monoclonal antibodies were able only partially to inhibit the binding of factor B to C3b indicates the presence of other binding site(s) within C3 for factor B. The finding that complete inhibition of binding could be achieved with the C3d fragment **(Fig. 4b)** may indicate that it contains more than one binding site for factor B.

Two anti-C3c monoclonal antibodies (F39b and F20b) were able to enhance the formation of the C3b-factor B complex. **(Table 1**). A possible explanation for the results observed could be that the monoclonal antibodies can stabilize the factor B-C3b complex resulting in tighter binding so that more factor B is bound to C3b at the end of the assay (i.e after washing steps).

Factor H was found partially to inhibit the binding of factor B to C3b **(Fig. 4b).** This indicates that factor H may bind to a site in C3b that is the same as or overlaps with one of the binding sites for factor B. This finding is consistent with previously reported data. DiScipio (1981) [9] and Kazatchkine et al. (1979) [15] reported that factor H was able to inhibit the binding of factor B to C3b-zymosan. A common binding site for factors B and H in C3d has been suggested by Koistinen et al (1989) [19] and Jokiranta (2001) [30].

The generation and purification of the three isolated domains of factor B made it possible to study their roles in binding to C3b **(Fig. 5b).** The expression of the vWF domain using an *E. coli* expression system [36] made it possible to study the role of this domain in the formation of the C3b-factor B complex. Competition binding studies were carried out separately with each of the individual domains as direct binding of ^125^ I-labelled domains/fragments could not be demonstrated. Presumably, binding of individual domains/fragments was weak and bound protein was removed during washing steps. It has previously been shown that isolated Ba or Bb fragments have no binding affinity for C3b [22, 23]. The results presented indicate that each of the three domains of factor B have binding sites for C3b as they could each inhibit binding by up to 50%. Direct or indirect evidence already exists for binding of each of the three domains of factor B to C3b. There is direct evidence for the role of the van Willebrand Factor domain in binding to C3b [44]. The Bb fragment was also able to inhibit binding of factor B to C3b by 50%. Similar inhibition was also seen in the presence of a mixture of Ba and Bb, suggesting that all the binding sites must be present in a single factor B molecule for complete inhibition to occur. The proposed two-site attachment model for the formation of the factor B-C3b complex [3, 21-24] is based upon evidence for single binding sites for C3b in each of the Ba and Bb fragments. There seems evidence for binding of each of the three domains of factor B to C3b.

In this system, the factor B bound to C3b-thiol Sepharose was susceptible to cleavage by factor D both in the presence of Mg^2+^ and EDTA **(Fig 6).** In the presence of either Mg^2+^ or EDTA, a large proportion of the factor B bound to C3b-thiol Sepharose was cleaved by factor D and maximum cleavage took place in a very short time (5 min). The Bb and Ba fragments were then released into the supernatant, consistent with the known short half-life of the C3bBb complex. Pryzdial and lsenman (1986) [12]) demonstrated that factor B present in their complex formed in the presence of EDTA in the fluid phase was also susceptible to cleavage by factor D.

## Abbreviations

CCP: complement control protein repeat
vWF: von Willebrand Factor
SP: serine protease
Pipes: [(Piperazine-N,N’-bis-2-ethanesulphonic acid). FPLC

## Acknowledgements

This research was funded by the Medical Research Council. SEW held an MRC Studentship.

